# The Role of Arthrobacter pascens 13LEP5 in mitigating Drought and Cold stress in Soybean (Glycine Max (L.) Merr.)

**DOI:** 10.64898/2026.08.03.742660

**Authors:** Yasha Jamil, Justina Kaziuniene, Giuseppe Colla, Sarune Ramoskaite, Monika Toleikiene

## Abstract

Drought and low temperatures are major abiotic factors affecting key physiological and biochemical processes and limiting the yields of soybean (Glycine max L. Merr.). To in-crease soybean production in Europe, different agricultural strategies are applied to re-duce abiotic stress, including biostimulants. Therefore, studies on the effectiveness of local strains isolated in Europe are becoming increasingly relevant. In this study two bacterial strains Arthrobacter pascens (AP) and Bradyrhizobium japonicum (BJ) along with plant-derived protein hydrolysate (PH) were analysed with soybean plans under abiotic stress conditions in plant growth chambers. Six treatments (control; AP; BJ; PH; BJ+AP; BJ+AP+PH) were tested to evaluate biostimulation effect before stress induction (VC stage) and to determine stress reduction effect on soybeans after plants recovery period (V3 stage). Biostimulants application has positive effect on soyabean biometric parameters in early plant development stage and post stress periods. More stable long-term effect was found on structural plant development parameters, than on pigment accumulation. The best results on plant biometric parameters were found where (AP) and (BJ+AP+PH) com-bination was inoculated. (BJ+AP+PH) combination was the only effective treatment, which showed significantly different results in pigments indices, compared to the control, after stress period.

**Author summary:** Yasha Jamil: Conceptualization, Data curation, Formal analysis, Writing– original draft, Giuseppe Colla: Formal analysis, Writing– original draft, Writing– review & editing, Justina Kaziuniene: Data curation, Formal analysis, Sarune Ramoskaite :Writing– review & editing. Monika Toleikienė: Conceptualization, Data curation, Formal analysis, Writing– original draft, Funding acquisition, Supervision, Writing– review & editing.

## Introduction

Soybean (Glycine max L. Merr.) is the world’s fourth most widely grown crop after corn, wheat and rice in terms of area harvested and production [1], [2]. Based on FAO-STAT data in 2024 the yearly output of soybean was over 397 million metric tons world-wide and only 17.12 million metric tons in Europe (EU) [3]. Soybean is nutritious plant, it contains the highest protein content (40%) and the second highest oil content (20%) among food legumes [4], [5]. Apart from proteins and oils, soybean contains basic nutritive con-stituents, such as vitamins, flavonoids, isoflavones, minerals, free sugar and saponins[6], [7]. Soybean is the only legume with alpha-linolenic acid and ample amount of essential omega-3 fatty acid [8], [9]. Soybeans are widely used in food and animal feed industry, it also used to produce vegetable oil, lecithin, polysaccharides and natural surfactants [10], [11]. According to the latest statistics, the value of soybeans and soybean products im-ported into Europe was 15.05 billion USD in 2022, 12.96 billion USD in 2023 and 11.8 bil-lion USD in 2024. The observed decline in the value of imported soybean and soybean products may be associated with the expanded soybean cultivation area in Europe. The harvested soybean area increased by 31.6%, between 2022 and 2024 [3]. However, such an increase does not meet the demand for soybean production, therefore, to reduce Europe’s dependence on imported soy products, it is important not only an expand of soybean cul-tivation area, but also to ensure a high-quality, abundant harvest.with alpha-linolenic acid and ample amount of essential omega-3 fatty acid [8], [9]. Soybeans are widely used in food and animal feed industry, it also used to produce vegetable oil, lecithin, polysaccharides and natural surfactants [10], [11]. According to the latest statistics, the value of soybeans and soybean products im-ported into Europe was 15.05 billion USD in 2022, 12.96 billion USD in 2023 and 11.8 bil-lion USD in 2024. The observed decline in the value of imported soybean and soybean products may be associated with the expanded soybean cultivation area in Europe. The harvested soybean area increased by 31.6%, between 2022 and 2024 [3]. However, such an increase does not meet the demand for soybean production, therefore, to reduce Europe’s dependence on imported soy products, it is important not only an expand of soybean cul-tivation area, but also to ensure a high-quality, abundant harvest.

Soybean is not domestic in Europe, this crop cultivation began in the late 19th century [12]. Due to the diverse soil and climatic conditions which are suboptimal in Europe, soy-bean cultivation still requires intensive research. Low air temperatures damage soybean production in northern Europe by suppressing soybeans growth in early growth stage, inducing flowers and pods abscission and insufficient grain filling at the pod-filling stage [13], [14]. Low soil temperatures disrupt root hydraulic conductance and reduce leaf tur-gor and expansion, while high irradiance can lead to photoinhibition [15]. Optimal tem-perature for soybean growth depends on the selected cultivar, latitude and growing tech-nology, but in general 22–32 °C temperature range is considered as the optimal and 17–18 °C temperature range is indicated as minimal temperature for soybean biological process-es. Air temperatures below 15 °C inhibit plant growth and development and temperatures below 10°C disrupt the flowering process [16], [17]. Drought stress is the most critical abi-otic stress which induces significant anatomical changes in soybean plants, impacting global soybean production, with approximately 40% of crop losses. Soil water deficit is typical problem for soybean cultivation in central and southern Europe [18], [19]. Water excess can also have negative impact for soybeans, hight moisture can increase pathogenic diseases development [17], [20], [21]. To improve soybean yields and grain quality differ-ent strategies are applied, including cultivar selection based on region climatic conditions [22], [23], pest and weed control [24], [25], [26], crop rotation [27], [28], tillage and water management [29], [30], fertilisation optimisation [29], [31] and plant stress reducing and growth biostimulating products application [32].

Research related with soybean abiotic stress reduction by using biostimulants are limited. Researchers are constantly searching for microorganisms or biologically active compounds that could reduce plant abiotic stress caused by drought and low tempera-tures. Biostimulants for abiotic stress reduction are classified into organic compounds (seaweeds, humic substances, amino acids, protein hydrolysates) and microorgan-isms-based (fungi, bacteria) products [33], [34]. Among all biostimulants, protein hydroly-sates are of particular interest, as it has positive effects on abiotic stress reduction. Protein hydrolysates contain phytohormones and amino acids that regulate action of signalling compounds responsible to stress management [35]. The major phytohormones responsible for reaction and regulation of abiotic stresses are abscisic acid (ABA), ethylene (ET), jasmonates (JA), salicylic acid (SA), cytokinins (CK), brassinosteroids (BR), auxins (AU), gibberellins (GA) and strigolactones (SLs) [36], [37]. ABA is a major stress phytohormone that plays a key role by regulating stress tolerance mechanisms [38], [39]. ABA promotes stomatal closure, stimulates cuticular wax formation, reduces mesophyll conductance, limits leaf expansion to minimize water loss. It also restricts shoot growth and lateral root development but promotes root elongation to improve water absorption [36], [40], [41]. Other phytohormones such ET, BR, SA, JA and SL are induced during abiotic stress and play critical roles in regulating plant defence response against abiotic stresses [36]. Other valuable compounds for abiotic stress reduction in the composition of protein hydroly-sates are amino acids. Amino acids such as proline, glycine betaine (GB), alanine, phenyl-alanine, aspartic acid, γ-aminobutyric acid (GABA), methionine, serine, cysteine, aspara-gine, glutamic acid, threonine, are well known for their ability to regulate stress signalling, antioxidative defence and osmoprotection [42], [43]. The rich composition of protein hy-drolysates and the lack of sufficient research on their effectiveness led us to conduct an experiment using plant protein hydrolysate to evaluate this biostimulation effect on soy-bean under abiotic stress conditions.

B. japonicum is one of the most significant bacteria for soybean plants. This symbiotic bacteria forms nodules on soybean roots and supply reduced atmospheric nitrogen (N2) for plant in ammonia form (NH3). In turn, the soybean provides carbon derived from photosynthesis and nutrients such as iron [44], [45]. Some research show that B. japonicum inoculants can not only increase soybean yields but also mitigate soil drought effect [46], [47], [48]. However, bradyrhizobia growth and persistence in soil may be also negatively impacted by drought conditions [49]. B. japonicum efficiency in adverse environment con-ditions can be improved by inoculating other bacteria additionally. It is important that strains selected for consortiums would be characterised as plant-growth-promoting rhi-zobacteria (PGPR) [46], [47]. Arthrobacter pascens is poorly studied species, described as halotolerant, Indole-3-Acetic Acid (IAA) producing, inorganic phosphorus dissolving and fluoranthene degrading bacteria [50]. A. pascens properties described in the scientific liter-ature, combined with the fact that this species has been successfully isolated from soy-beans, suggesting that this bacterium may have potential as a biostimulant for soybean growth promotion and even abiotic stress reduction.

In this research, we will investigate A. pascens 13LEP5, B. japonicum, plant-derived protein hydrolysate and their combinations efficiency on soybean drought and cold caused stress reduction. We will analyse biostimulants effect on soybean biometric and spectral parameters before (VC growth stage) and after (V3 growth stage) abiotic stress in-duction. This study provides the first assessment of A. pascens in association crop plants (soybean), providing a direct evaluation of its biostimulatory potential and its ability to reduce abiotic stress.

## Materials and methods

### 2.1. Origin of Biostimulants

Three different biostimulants were selected for this research to evaluate their potential for abiotic stress reduction. Commercially available inoculant “BACTOLiVE®LEGUME” (Rhizo-mic GmbH, St. Johann, Germany) suitable for different legume pants growth pro-motion, including soybean, was selected for this experiment as a reliable and effective product on the market. This product contains B. japonicum and other rhizobia strains.

The A. pascens 13LEP5 strain was obtained from the Lithuanian Research Centre for Agri-culture and Forestry microorganism collection. This strain was isolated in 2023, from soy-bean seedlings leaf’s surface. Soybean seedlings were collected in Lithuania, Kėdainiai distr., Dotnuva [55°24′ N, 23°51′ E], before blooming at fifth–sixth trifoliolate formation growth stage (between V–5 and V–6). A. pascens 13LEP5 strain was grown in Luria Broth (LB) [Tryptone 10 gL-1 (Organotechnie S.A.S., La Courneuve, France), 5 gL-1 Yeast Extract (Organotechnie S.A.S., La Courneuve, France), 10 gL-1 NaCl (Sigma-Aldrich, St. Louis, MO, USA)][51], for 48 hours at 30°C temperature, 130 rpm. Cell count of final bacterial suspen-sion was performed based on serial dilution method, using the same LB medium supple-mented with 20 gL-1 agar (SERVA Electrophoresis GmbH, Germany). Cell count of final A. pascens 13LEP5 suspension was 1.0×107 CFUmL-1.

Plant-derived protein hydrolysate “Trainer” was obtained from University of Tuscia, Italy. Based on product composition analysis this protein hydrolysate contains approxi-mately 35.5% of organic matter, 5% of total nitrogen, and 27% of amino acids and soluble peptides [52].

### 2.2. Molecular identification

13LEP5 strain identification was carried out using partial 16S rDNA sequence analy-sis. All partial 16S rDNA sequences were determined by PCR with primers 8 F (5′-AGAGTTTGATCCTGGCTCAG-3′) and 1492R (5′-GGTTACCTTGTTACGACTT-3′) [53] and compared to the EzBiocloud identification service. Phylogenetic tree was constructed by comparing obtained 16S rDNA sequences with related type strains 16S rDNA se-quences. Phylogenetic tree was constructed using MEGA 5.0 software [54].

### 2.3. Soybean Growth Experiment Modulating Abiotic Stress Conditions

Soybean seeds were sterilised with 70% ethyl alcohol solution, then washed with sterile deionized water (SDW) and additionally disinfected with sodium hypochlorite so-lution containing 5% active chlorine for 5 min. Finally, soybean seeds were rinsed 5 times with SDW [55]. Sterile seeds were coated with different products based on (Table 1).

**Table 1.** Soybean seed coating scheme and growth conditions.

| **CODE** | **TREATMENT** | **SOYBEAN GROWTH CONDITIONS** |  |  |
| --- | --- | --- | --- | --- |
|  |  | **Optimal period**<br>**(VE-VC)** | **Stress induction**<br>**(VC-V1)** | **Recovery period**<br>**(V1 - V3)** |
| *Control* | Water | 16/8-h day/night photoperiod, 20/18 °C day/night temperature mode, irrigation with DSW 80 mL/pot/day | 16/8-h day/night pho-to-period, I and VII days - 20/18 °C, II and VI days - 14/13°C, III and V days - 8/7°C, IV day - 5/4°C day/night temperature mode, irrigation with DSW 35 mL/pot/day | 16/8-h day/night pho-to-period, 20/18 °C day/night temperature mode, irrigation with DSW 100-150 mL/pot/day. |
| *AP* | A. pascens 13LEP5 |  |  |  |
| *BJ* | B. japonicum |  |  |  |
| *PH* | Protein hydrolysate |  |  |  |
| *BJ+AP* | A combination of B. japonicum and A. pascens 13LEP5 |  |  |  |
| *(AP+BJ+PH)* |  |  |  |  |

Each treatment was prepared by using 10 mL of product or products combination solution for 10 g soybean seeds coating. A suspension of the peat-based powder “BAC-TOLiVE®LEGUME” containing B. japonicum was prepared by mixing, 0.04 g powder with 1 ml DSW and shaken for 5 min to detach bacteria cells from the peat. Prepared bacterial suspension was diluted with DSW to a final volume of 10 mL. A. pascens 13LEP5 suspen-sion was diluted with DSW (ratio 1:1). The plant-derived protein hydrolysate treatment was prepared by diluting 200 µL of the product with 9.8 mL of DSW. Combinations of products were prepared using the undiluted components, which were subsequently mixed and adjusted to a final volume of 10 mL with DSW. Coated soybean seed were sown in 10 cm diameter pots filled with peat moos, vermiculite and quartz sand mix in a ratio (2:1:1). Four seeds were sown in pot. Each variant was grown in four repetitions.

Soybean was grown in the plant growth chamber in optimal growth conditions 16/8-h day/night photoperiod and 20/16 °C day/night temperature modes and optimal water rate of 80 mL/day/pot was applied until seedlings reached cotyledon stage (VC). During VC growing period lower temperature and daily water application were adjusted in growth chamber to induce abiotic stress for soybean seedlings. Temperature modes were adjusted every day for week (Table 2), watering rate was 35 mL/day/pot.

**Table 2.** Temperature modes during soybean stress induction period.

| Time | Temperature °C ranges, in different days |  |  |  |  |  |  |
| --- | --- | --- | --- | --- | --- | --- | --- |
|  | I | II | III | IV | V | VI | VII |
| Day | 20 | 14 | 8 | 5 | 8 | 14 | 20 |
| Night | 18 | 13 | 7 | 4 | 7 | 13 | 18 |

After stress induction period 20/16 °C day/night temperature modes and optimal wa-ter rate of 100-130 mL/day/pot were adjusted. Soybeans seedlings were grown until V3 growth stage.

**Table 3.**
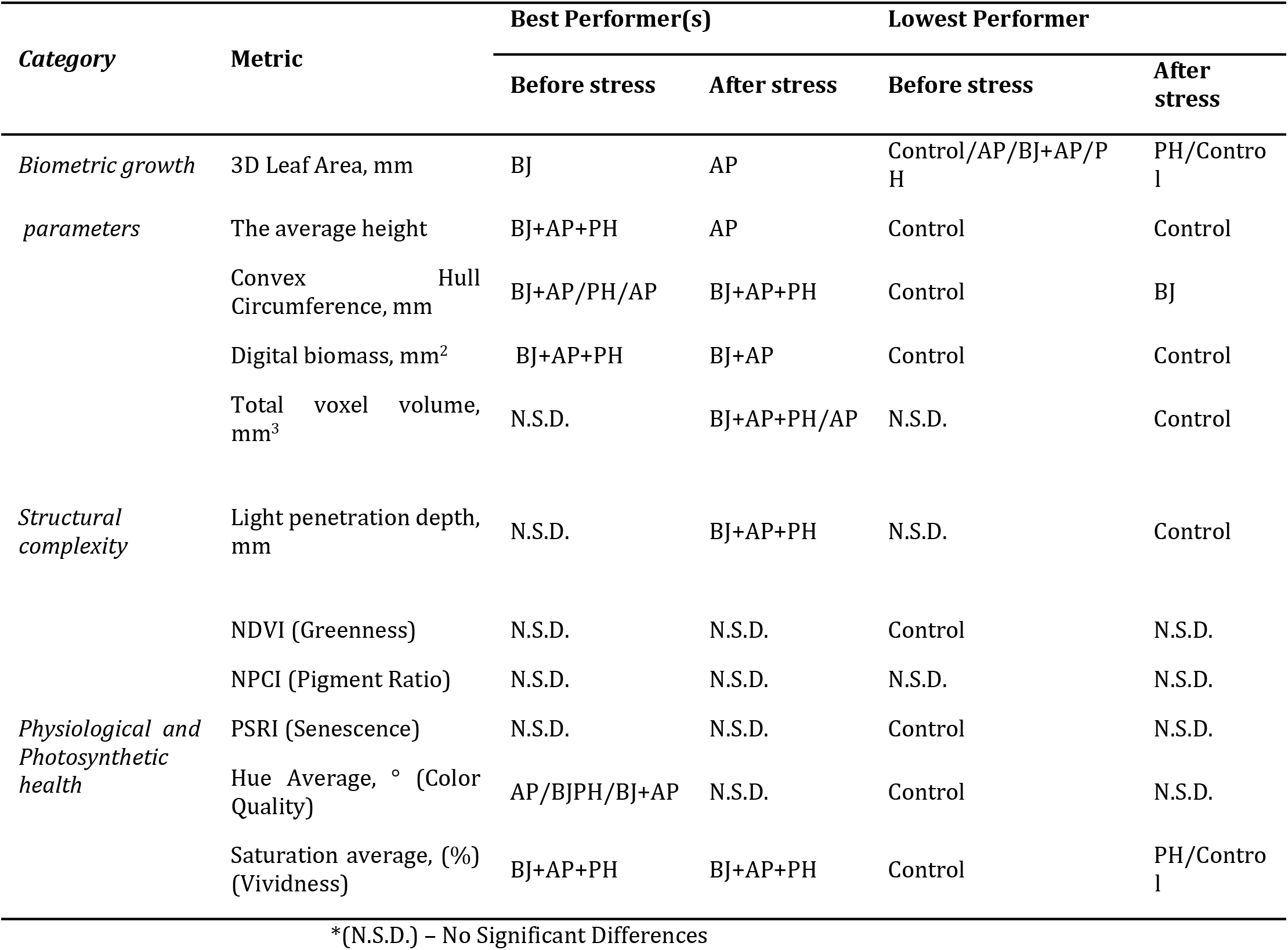
Biostimulants efficiency on soybean growth parameters. Soybean treatments: A. pascens 13LEP5 (AP), B. japonicum (BJ), protein hydrolysate (PH), combination of B. japonicum and A. pascens 13LEP5 (BJ+AP), combination of B. japonicum, A. pascens 13LEP5 and protein hydrolysate (BJ+AP+PH).

### 2.4. Phenotyping of soybean above part

Morphological, structural, and physiological characteristics of soybean seedlings were analysed two times in VC (before stress induction) and V3 (recovery period) growth stages. 3D Leaf Area (mm), the average height (mm), convex hull circumference (mm), digital bio-mass (mm2), total voxel volume (mm3), light penetration depth (mm), NDVI, NPCI, PSRI, Hue Average and saturation average (%) were measured to describe the canopy growth, light penetration and the response to the stress by using high-resolution 3D multispectral scanner PlantEye F500(Phenospex, Heerlen, The Netherlands) [56]. All data were auto-matically computed by Hortcontrol v. 3.8 software [56], [57].

### 2.5. Statistical Analysis

Outliers were verified by interquartile range and mean with standard deviations were calculated in every treatment and parameter. The statistical analysis was performed with the R software (version 4.3.2) [58]. The effect of treatments on the morphological and spec-tral characteristics was analysed by one-way analysis of variance (ANOVA) and the sig-nificant differences between the means were found using Duncans Multiple Range Test (DMRT) [59]. The smallest significant difference was calculated using a probability level of p ≥ 0.05. The ggplot2 package was used to create graphical visualization and letters (a, b, c) were used to indicate statistically significant differences between the treatments[60]. The values marked with the same letter showed no significant difference at p ≥ 0.05.

## 3. Results

### 3.1. 16S rRNA phylogenetic tree

13LEP5 isolate 16S rRNA sequence were compared with the 16S rRNA gene sequences of related type strains and phylogenetic tree was constructed (Fig. 1). Phylogeneetic tree showed that 13LEP5 isolate belongs to Arthrobacter spp. and is closely related to Arthrobac-ter pascens species. 13LEP5 and Arthrobacter pascens DSM 20545 strains were homologous with percent identity 99.57 %.

**Figure 1.**
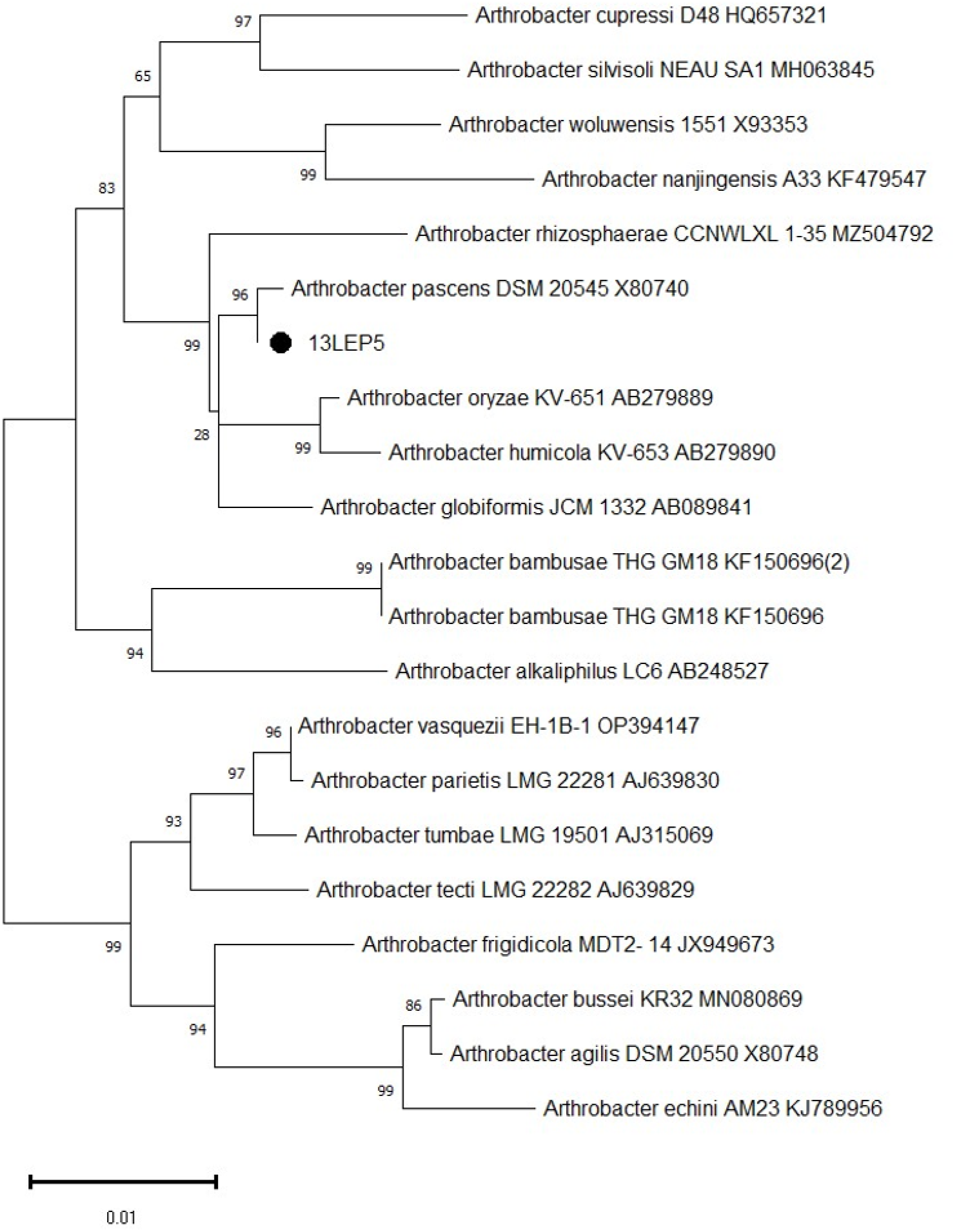
The phylogenetic relationships of 13LEP5 isolate within the genus of Arthrobacter spp. investigated using 16S rRNA gene sequence analysis. The phylogenetic tree was constructed by using MEGA 5.0 software package, neighbour-joining statistical method with 1000 Bootstrap rep-licates. The scale bar illustrates 0.01 substitutions per nucleotide position.

### 3.2. Biometrical Characteristics of Soybean

Different parameters were analysed to evaluate abiotic stress influence on soybean seed-lings growth, architecture, light interaction, physiological and photosynthetic health. Re-sults demonstrated that different biostimulants had different efficiency on soybean seed-lings stress reduction.

The 3D leaf area of soybean plants varied significantly across treatments both before the induction of environmental stress and following the recovery period (Figure 2). Prior to the application of drought and cold stress (VC stage), the seed treatment with (B) resulted in the highest 3D leaf area (approximately 3419 mm2), significantly outperforming the control (approximately 1323 mm2), and other individual treatments, except (BJ+AP+PH) variant (p<0,05). Following the seven-day stress period and subsequent three-week recov-ery, a shift in treatment efficacy was observed. Plants treated with (AP) exhibited the most robust recovery, achieving the highest numerical 3D leaf area (approximately 3905 mm2), which was significantly greater than control (approximately 2033 mm2) and plant-derived protein hydrolysate variant (approximately 1742 mm2).

**Figure 2.**
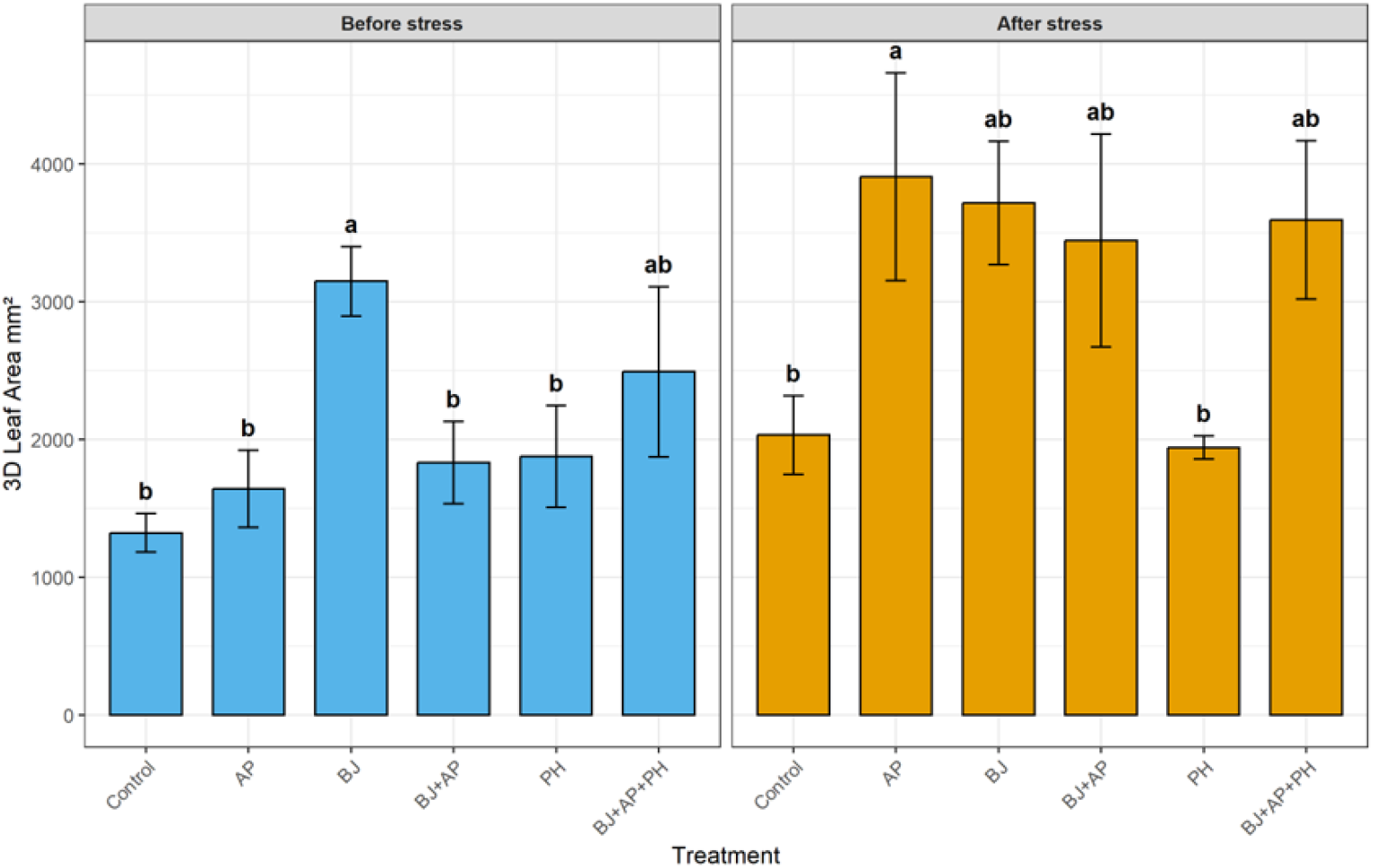
Leaf area (mm2) of soybean seedlings, following treatment with different products, before (VC growth stage) and after (V3 growth stage) stress induction. Soybean treatments: A. pascens 13LEP5 (AP), B. japonicum (BJ), protein hydrolysate (PH), combination of B. japonicum and A. pascens 13LEP5 (BJ+AP), combination of B. japonicum, A. pascens 13LEP5 and protein hydrolysate (BJ+AP+PH). Error bars indicate the standard deviations within biological replications at each treatment. Values marked with the same letter are not significantly different at p ≥ 0.05.

Plant height was significantly improved almost by all biostimulants applications compared to the control across both measurement periods (Figure 3). In the pre-stress phase, the triple combination (BJ+AP+PH) promoted the greatest vertical elongation and was significantly higher than control 28 mm and (AP) variant 36 mm, achieving a height of approximately 54 mm. After recovery phase the tallest plants were obtained where (AP) strain was applied, average height was 68 mm and significantly deferred from control, (BJ) and (BJ+AP+PH) biostimulants treated plants where average plant height was 47 mm, 52 mm and 54 mm respectively. Conversely, the (BJ+AP+PH) treatment, which initially led in height, showed a comparatively lowest growth rate during the recovery stage, finishing in statistical group “bc.“

**Figure 3.**
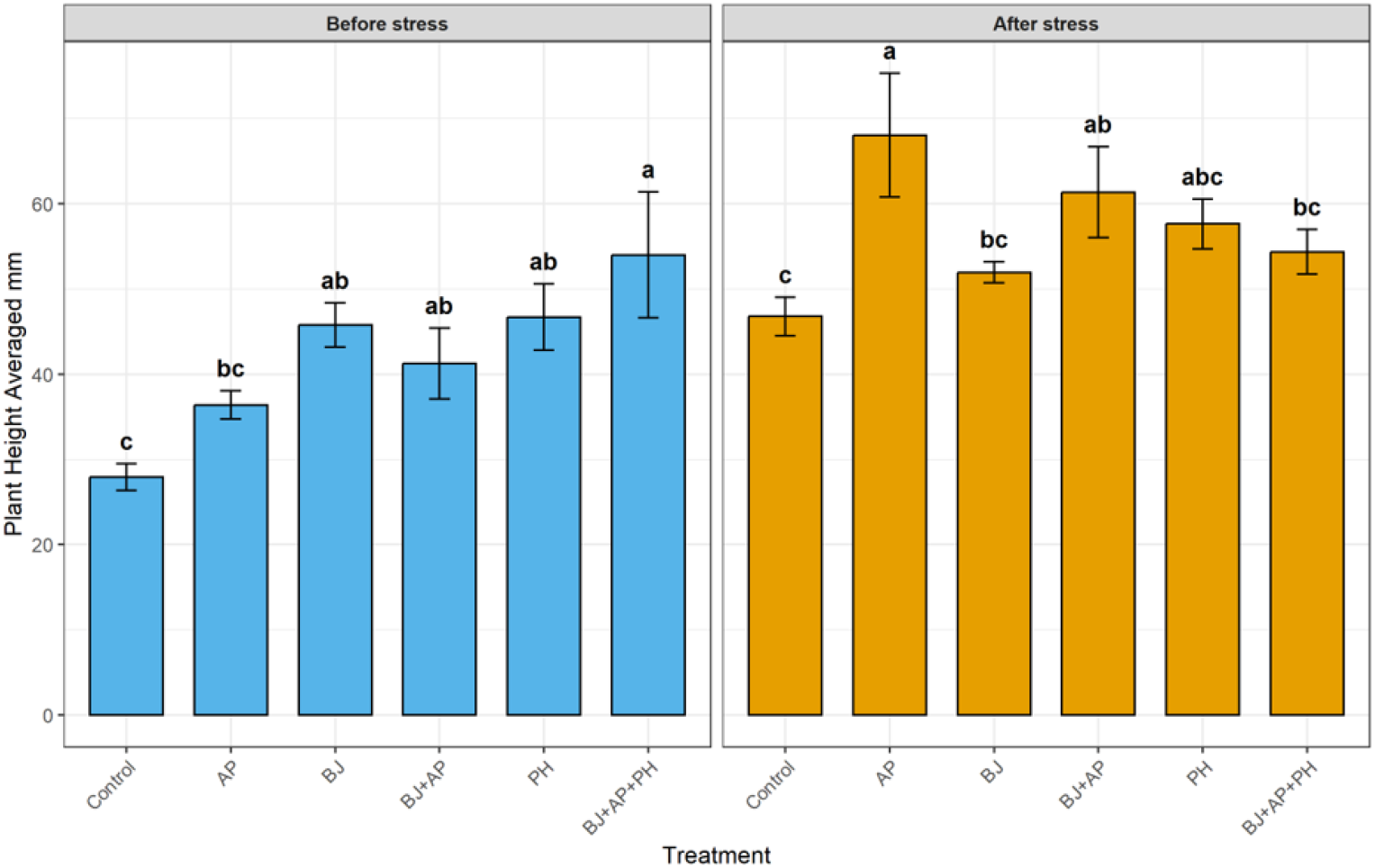
The average height (mm) of soybean seedlings, following treatment with different products, before (VC growth stage) and after (V3 growth stage) stress induction. Soybean treat-ments: A. pascens 13LEP5 (AP), B. japonicum (BJ), protein hydrolysate (PH), combination of B. ja-ponicum and A. pascens 13LEP5 (BJ+AP), combination of B. japonicum, A. pascens 13LEP5 and protein hydrolysate (BJ+AP+PH). Error bars indicate the standard deviations within biological replications at each treatment. Values marked with the same letter are not significantly different at p ≥ 0.05.

The Convex Hull Circumference (mm), which quantifies the horizontal perimeter of the plant’s spatial footprint, was significantly influenced by treatment application (Figure 4). Before the induction of stress, the (BJ+AP) dual microbial treatment promoted the great-est spatial spread 326 mm (Group a) and significantly differed only from control and (BJ) variants where horizontal perimeter was 154 mm and 182 mm. While the most biostimu-lants treatments significantly increased circumference compared to the control, BJ alone resulted in a more compact, significantly not different architecture, compared to the un-treated variant.

**Figure 4.**
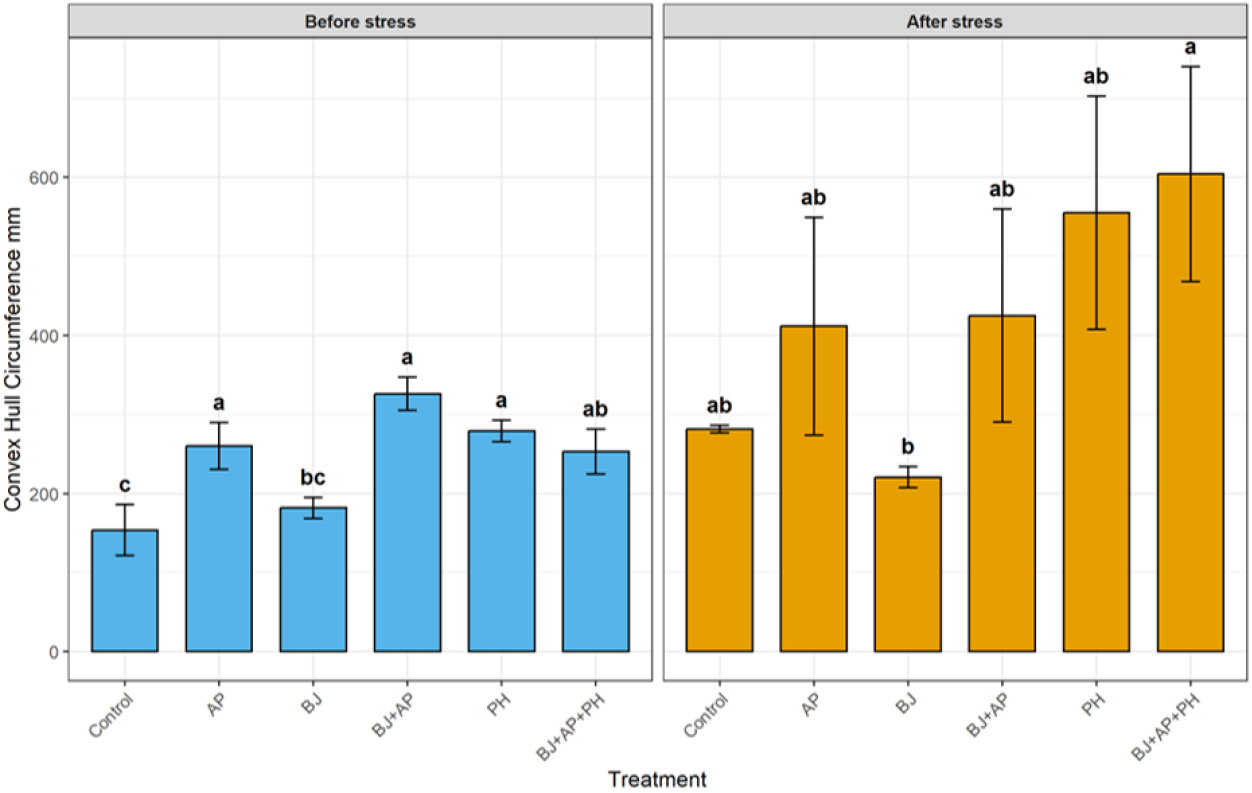
Convex hull circumference (mm) of soybean seedlings, following treatment with different products, before (VC growth stage) and after (V3 growth stage) stress induction. Soybean treat-ments: A. pascens 13LEP5 (AP), B. japonicum (BJ), protein hydrolysate (PH), combination of B. ja-ponicum and A. pascens 13LEP5 (BJ+AP), combination of B. japonicum,A. pascens 13LEP5 and protein hydrolysate (BJ+AP+PH). Error bars indicate the standard deviations within biological replications at each treatment. Values marked with the same letter are not significantly different at p ≥ 0.05.

Following the recovery phase, a highest expansion was observed in the triple combi-nation (BJ+AP+PH). The (BJ+AP+PH) combination achieved the highest circumference 604 mm (Group a), indicating a robust horizontal expansion during recovery, however, result obtained was significantly higher only compared with (BJ), where horizontal perimeter of the soybean was 220 mm, no significant differences were obtained compared with other variants. Conversely, the (BJ) treatment remained the most compact among the biostimu-lants (Group b).

The digital biomass was significantly influenced by the biostimulant treatments throughout the experimental period (Figure 5). Prior to the induction of stress, almost all biostimulants, excluding (BJ), significantly increased digital biomass of soybean seedlings, compared to the control variant. The application of (AP), (BJ+AP) and the triple combina-tion of (BJ+AP+PH) resulted in the highest biomass accumulation 63482 mm3, 53365 mm3 and 51455 mm3 respectively, showing a significant increase compared to the control and (BJ) variants, where digital biomass accumulated was 21015 mm3 and 21996 mm3. Fol-lowing the recovery phase, only (BJ+AP) treated plants maintained significantly higher digital biomass 58083 mm3, compared to the control 29274 mm3.

**Figure 5.**
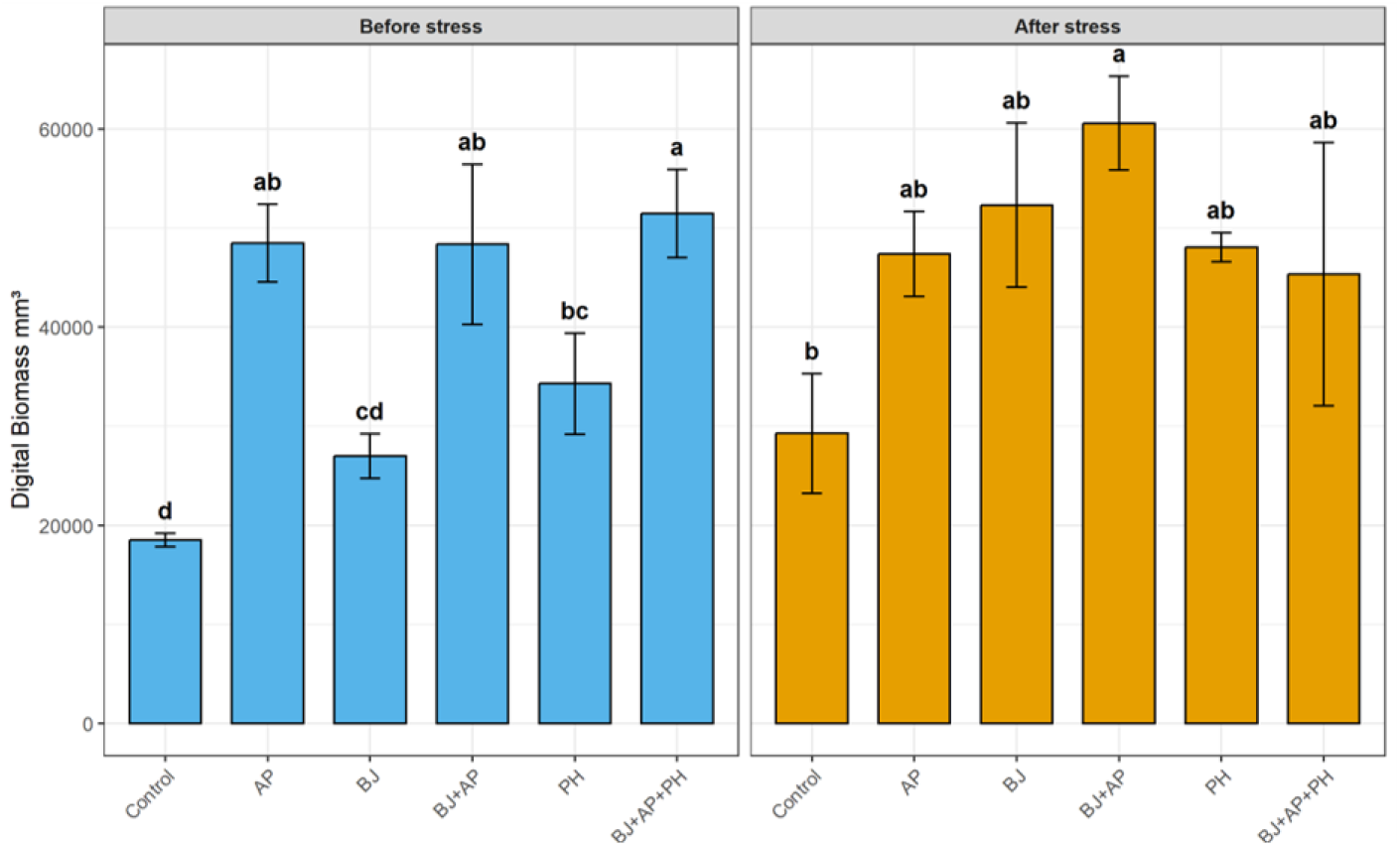
Digital biomass (mm2) of soybean seedlings, following treatment with different products, before (VC growth stage) and after (V3 growth stage) stress induction. Soybean treatments: A. pascens 13LEP5 (AP), B. japonicum (BJ), protein hydrolysate (PH), combination of B. japonicum and A. pascens 13LEP5 (BJ+AP), combination of B. japonicum, A. pascens 13LEP5 and protein hydrolysate (BJ+AP+PH). Error bars indicate the standard deviations within biological replications at each treatment. Values marked with the same letter are not significantly different at p ≥ 0.05.

The total voxel volume, representing the cumulative 3D space occupied by plant tis-sue, showed a highly significant response to biostimulant application following abiotic stress (Figure 6). During the initial growth phase, no statistical differences were observed between treatments (Group a). However, following the recovery phase, the application of (AP) and the triple combination (BJ+AP+PH) resulted in the significantly highest voxel volumes 5846 mm3 and 5973 mm3 (Group a) and was significantly higher compared to control, (BJ) and (PH) variants, where total voxel volume was 3810 mm3, 4744 mm3 and 4544 mm3, respectively. Single application with (BJ) (Group b) and dual microbial combi-nation (BJ+AP) (Group ab) also demonstrated significantly better robust volume recovery compared to the untreated control plants (Group c). No significant differences were ob-tained between the (PH) (Group bc) and control (Group c) total voxel volume results.

**Figure 6.**
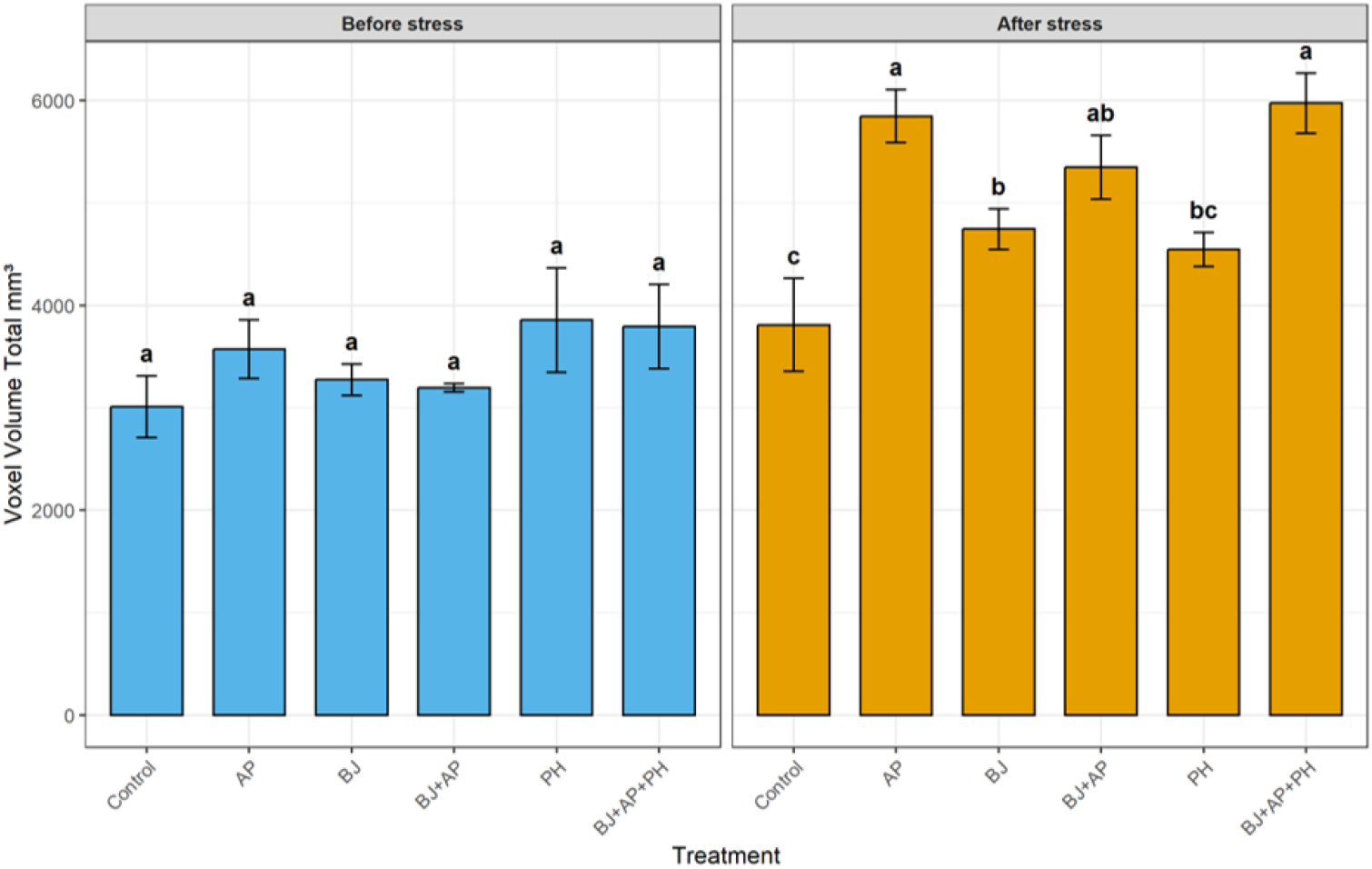
Total voxel volume (mm3) of soybean seedlings, following treatment with different products, before (VC growth stage) and after (V3 growth stage) stress induction. Soybean treat-ments: A. pascens 13LEP5 (AP), B. japonicum (BJ), protein hydrolysate (PH), combination of B. ja-ponicum and A. pascens 13LEP5 (BJ+AP), combination of B. japonicum, A. pascens 13LEP5 and protein hydrolysate (BJ+AP+PH). Error bars indicate the standard deviations within biological replications at each treatment. Values marked with the same letter are not significantly different at p ≥ 0.05.

### 3.3. Structural complexity of the soybean canopy

The structural complexity of the soybean canopy was further evaluated through light penetration depth (mm). During the pre-stress phase, no significant differences were observed between treatments, with all groups exhibiting a similar light penetration depth (Figure 7). Following the recovery period, the (BJ+AP+PH) triple combination resulted in the highest light penetration depth 50 mm (Group a), significantly higher compared to the control plants, where light penetration was 36 mm (Group b), but with no statistical dif-ferences compared with other biostimulats treated plants (Group ab).

**Figure 7.**
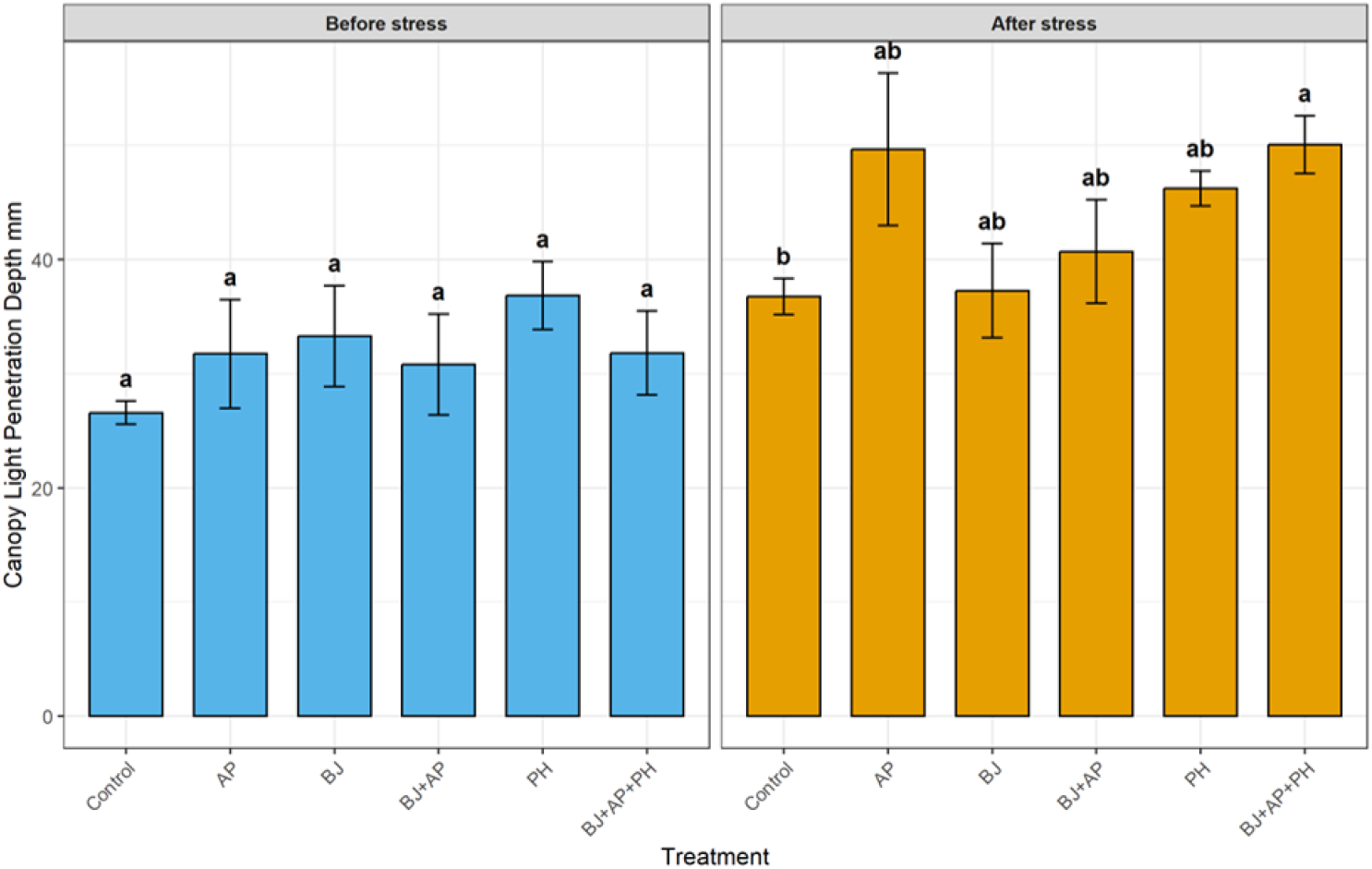
Light penetration depth (mm) in soybean seedlings, following treatment with different products, before (VC growth stage) and after (V3 growth stage) stress induction. Soybean treat-ments: A. pascens 13LEP5 (AP), B. japonicum (BJ), protein hydrolysate (PH), combination of B. ja-ponicum and A. pascens 13LEP5 (BJ+AP), combination of B. japonicum, A. pascens 13LEP5 and protein hydrolysate (BJ+AP+PH). Error bars indicate the standard deviations within biological replications at each treatment. Values marked with the same letter are not significantly different at p ≥ 0.05.

### 3.4. Physiological and Photosynthetic health

The Normalized Difference Vegetation Index (NDVI) was used as a metric for chlo-rophyll content and overall photosynthetic health. Prior to the stress period, all biostimu-lant-treated plants exhibited significantly higher NDVI values (ranging from 0.58 to 0.62) compared to the control variant 0.48, indicating enhanced early-stage physiological vigor across all treated groups (Figure 8). Following the seven-day stress period and the subse-quent three-week recovery phase, the statistical differences between treatments vanished. All experimental groups, including the control, reached a similar physiological state (Group a), with NDVI values stabilizing between 0.53 and 0.62.

**Figure 8.**
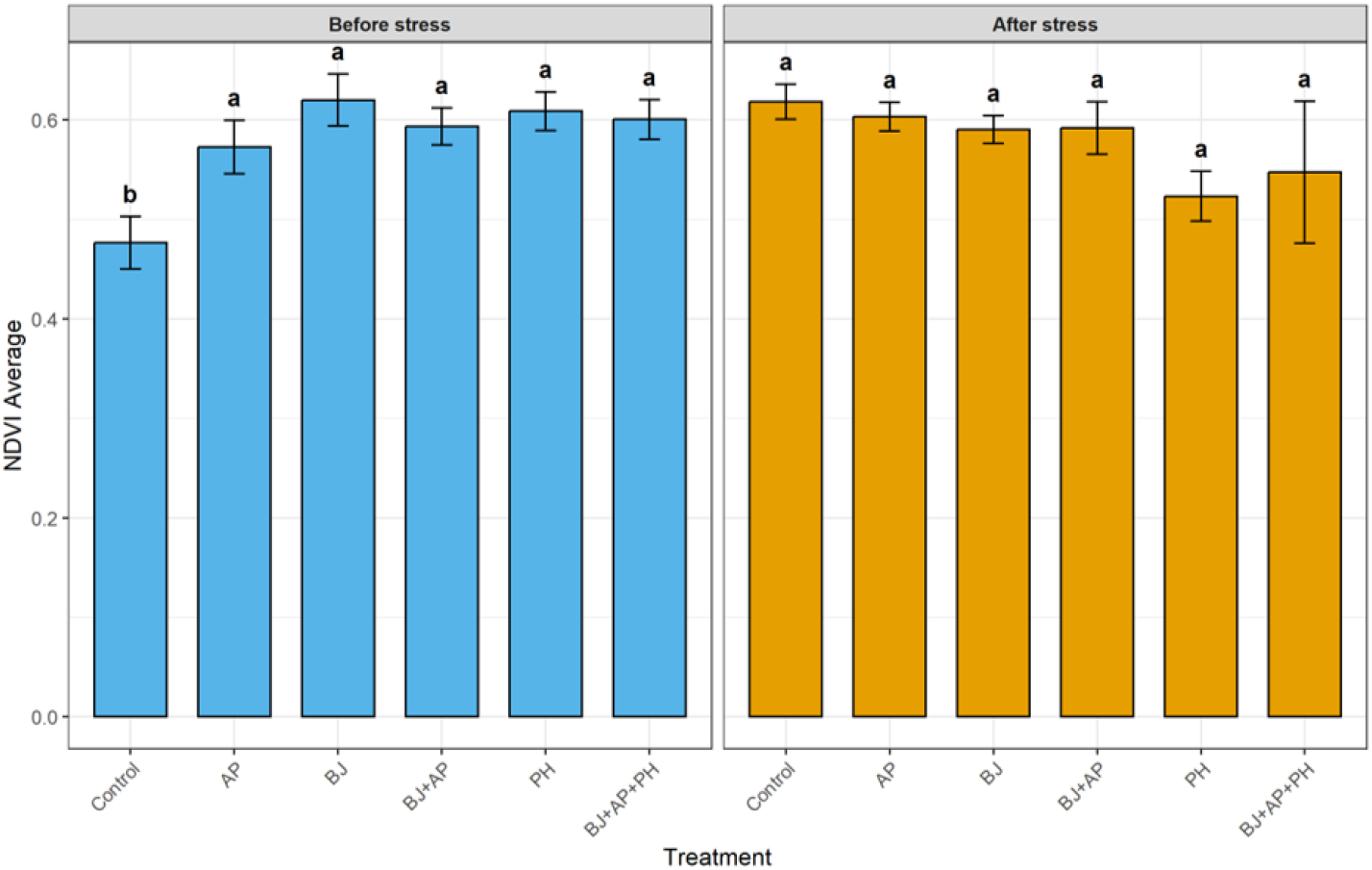
The Normalized Difference Vegetation Index (NDVI) in soybean seedlings, following treatment with different products, before (VC growth stage) and after (V3 growth stage) stress induction. Soybean treatments: A. pascens 13LEP5 (AP), B. japonicum (BJ), protein hydrolysate (PH), combination of B. japonicum and A. pascens 13LEP5 (BJ+AP), combination of B. japonicum, A. pascens 13LEP5 and protein hydrolysate (BJ+AP+PH). Error bars indicate the standard deviations within biological replications at each treatment. Values marked with the same letter are not significantly different at p ≥ 0.05.

The Normalized Pigment Chlorophyll Index (NPCI), did not showed any significant sensitivity to the biostimulant treatments under the pre-stress and after stress periods (Figure 9). In both periods all treatments, including control, showed significantly not dif-ferent (NPCI) results, ranging from 0.04 to 0.08 (Group a).

**Figure 9.**
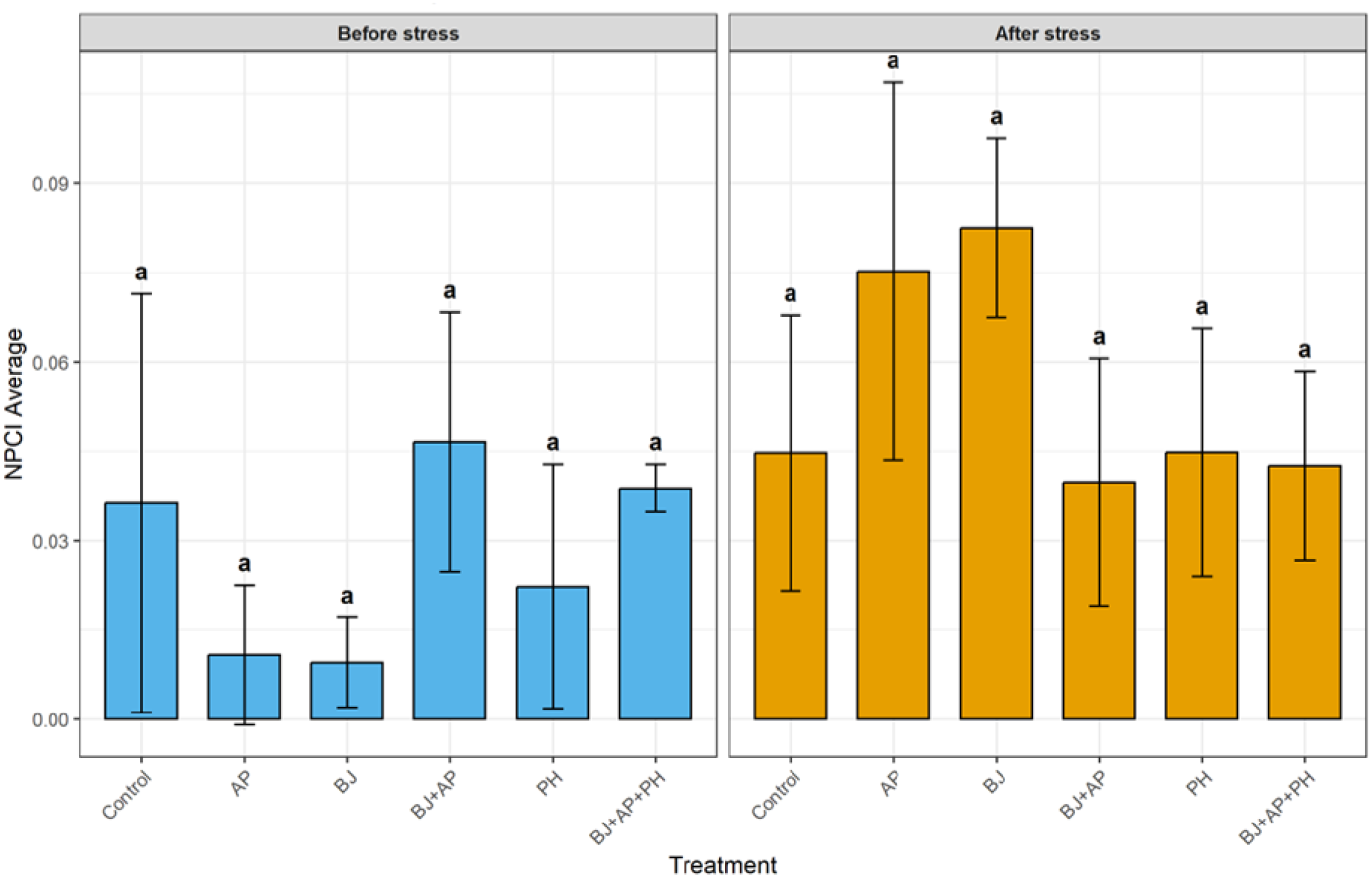
The Normalized Pigment Chlorophyll Index (NPCI) in soybean seedling, following treatment with different products, before (VC growth stage) and after (V3 growth stage) stress induction. Soybean treatments: A. pascens 13LEP5 (AP), B. japonicum (BJ), protein hydrolysate (PH), combination of B. japonicum and A. pascens 13LEP5 (BJ+AP), combination of B. japonicum, A. pascens 13LEP5 and protein hydrolysate (BJ+AP+PH). Error bars indicate the standard deviations within biological replications at each treatment. Values marked with the same letter are not significantly different at p ≥ 0.05.

The Plant Senescence Reflectance Index (PSRI) was assessed to determine the degree of leaf senescence and physiological stress. Significant differences were observed during the pre-stress phase (Figure 10). The untreated control plants exhibited the highest PSRI value 0.05 (Group a). In contrast, all biostimulant treatments, particularly (BJ), maintained significantly lower PSRI values (Group b). Following the recovery period, the PSRI values across all treatments stabilized, with no statistically significant differences remaining (Group a).

**Figure 10.**
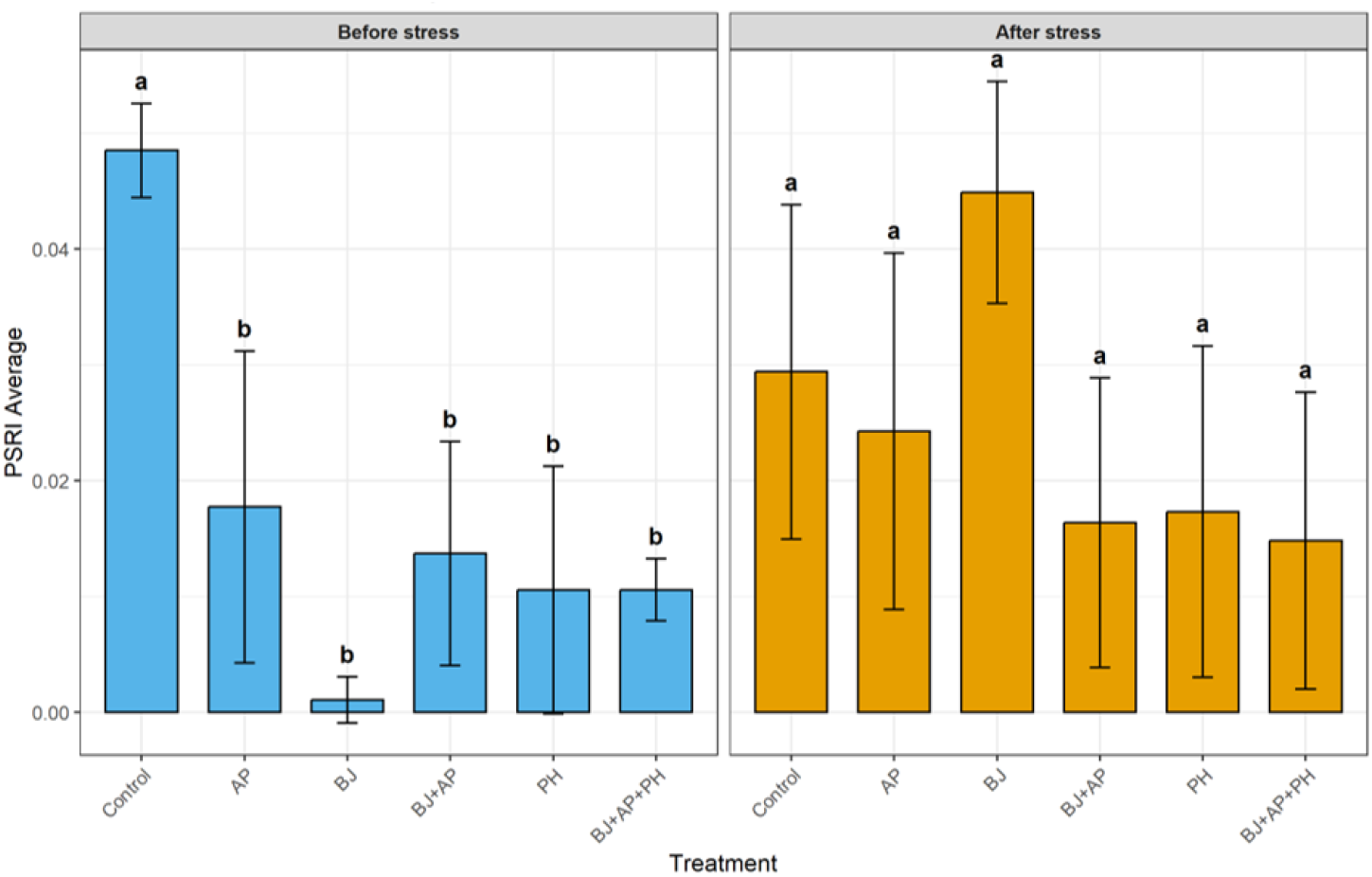
The Plant Senescence Reflectance Index (PSRI) in soybean seedling, following treatment with different products, before (VC growth stage) and after (V3 growth stage) stress induction. Soybean treatments: A. pascens 13LEP5 (AP), B. japonicum (BJ), protein hydrolysate (PH), combina-tion of B. japonicum and A. pascens 13LEP5 (BJ+AP), combination of B. japonicum, A. pascens 13LEP5 and protein hydrolysate (BJ+AP+PH). Error bars indicate the standard deviations within biological replications at each treatment. Values marked with the same letter are not significantly different at p ≥ 0.05.

The Hue Average (°) was analysed to quantify the colour shade of the soybean cano-py. Prior to the induction of stress, all biostimulant treatments significantly enhanced the leaf hue compared to the control (Figure 11). The applications of (AP), (BJ), (BJ+AP) and (PH) resulted in a more vibrant green canopy 119.5°, 118.5° and 116.0° respectively (Group a) compared to the control plants (Group b). Following the recovery period, the hue values across all treatments stabilized and showed no significant differences.

**Figure 11.**
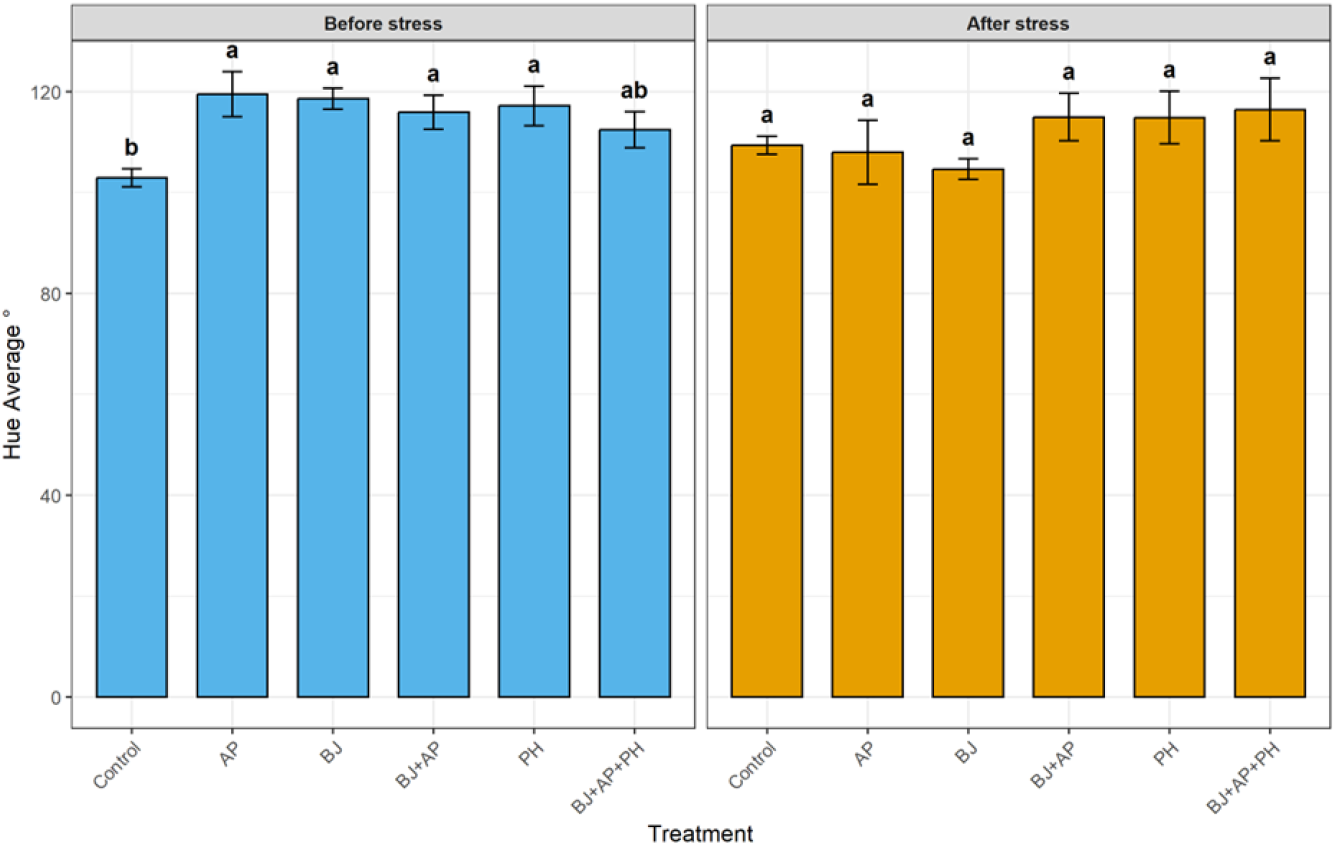
The hue average in soybean seedling, following treatment with different products, before (VC growth stage) and after (V3 growth stage) stress induction. Soybean treatments: A. pascens 13LEP5 (AP), B. japonicum (BJ), protein hydrolysate (PH), combination of B. japonicum and A. pascens 13LEP5 (BJ+AP), combination of B. japonicum, A. pascens 13LEP5 and protein hydrolysate (BJ+AP+PH). Error bars indicate the standard deviations within biological replications at each treatment. Values marked with the same letter are not significantly different at p ≥ 0.05.

The saturation average (%), representing the intensity and vividness of leaf colora-tion, was significantly influenced by the biostimulant treatments (Figure 12). Prior to stress, the triple combination (BJ+AP+PH) produced the highest saturation 28.9 % (Group a) and was significantly higher compared to the control variant 24.9 %, while all other bi-ostimulants insignificantly surpassed control variant (Group b). Following the recovery phase, the (BJ+AP+PH) treatment achieved the highest saturation value 33.5 % (Group a) and was significantly higher that control 27.4 % and (PH) 25.4 % variants. (PH) treatment alone resulted the lowest saturation (Group c).

**Figure 12.**
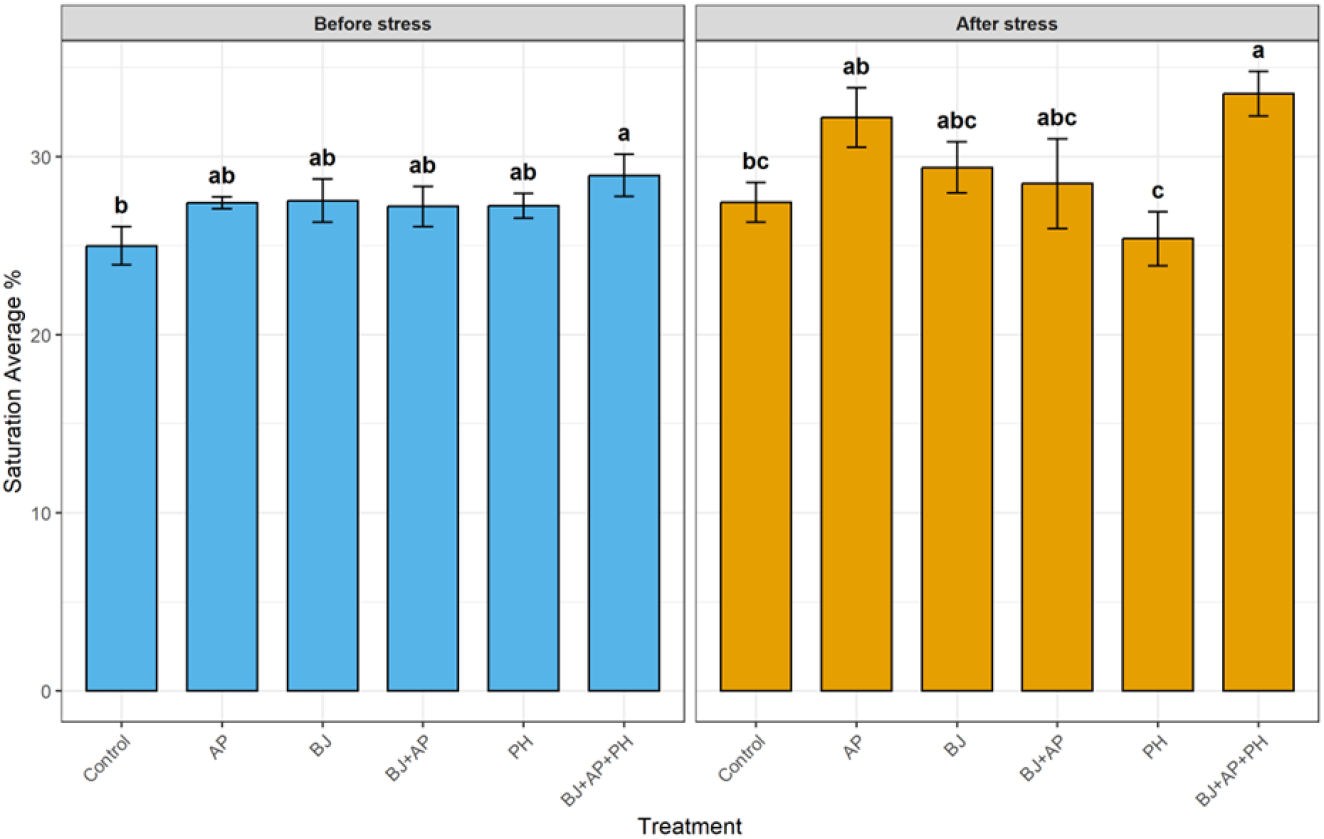
The saturation average (%) in soybean seedling, following treatment with different products, before (VC growth stage) and after (V3 growth stage) stress induction. Soybean treat-ments: A. pascens 13LEP5 (AP), B. japonicum (BJ), protein hydrolysate (PH), combination of B. ja-ponicum and A. pascens 13LEP5 (BJ+AP), combination of B. japonicum, A. pascens 13LEP5 and protein hydrolysate (BJ+AP+PH). Error bars indicate the standard deviations within biological replications at each treatment. Values marked with the same letter are not significantly different at p ≥ 0.05.

### 3.4. Biostimulants effeciency on soybean growth under abiotic stress conditions

Different biostimulants had different efficiency on soybean growth, structural, physi-ological and health parameters before and after modulated drought and cold stress condi-tions. The best results before stress induction were obtained with single B. japonicum (BJ) inoculation and combinations where B. japonicum (BJ) included. After stress induction and recovery period the best performance were obtained with single application of A. pascens 13LEP5 (AP) strain and triple combination of (BJ+AP+PH). B. japonicum (BJ) incorporation into combinations with other biostimulants provided significantly more value for stress affected soya seedlings, compared to single (BJ) applications. Protein hydrolysate (PH) has low efficiency in single applications before and after stress reduction and demonstrated average results.

## 4. Discussion

Our results showed that biostimulants combinations are more affective on plant bi-ostimulation than single applications. Although biostimulants synergistic effects are less studied, several scientific research show that biostimulants combinations are more benefi-cial for plant than single biostimulants applications. Savarese and her colleagues found that plants treated with potassium humates and green compost tea combination (MIX) accumulated significantly larger biomass production and nutrient uptake compared to the control and single potassium humate (KH) or green compost tea (CT) treated plants. They also found that MIX and microbial product combination (MIX_M+) significantly increased shoot dry weight by 52 %, 30 %, 31 %, 31 %, 13%, 24 %, 21 % compared to control, (M+), (KH), (CT), (MIX), (KH_M+) and (CT_M+) treatments, respectively [61]. Mishra ant his col-leagues performed biostimulants combinations efficiency investigations on maize under drought conditions. It was revealed that the combination of Adhatoda vasica leaf extract and Pseudomonas putida significantly increased root length, shoot length, plant fresh weight, and dry weight by approximately 7.7%, 21.1%, 24.2%, and 42.7%, respectively, compared to the single application of A. vasica. The same combination significantly increased the root length, shoot length, plant fresh weight and dry weight by 15.5 %, 40.8 %, 40.7 %, 46.8 %, respectively, compared to single P. putida application [62]. Our investigation results con-firmed other scientist statements that biosimulants combinations are more affective on plant biostimulation than single applications. Additionally, our results suggest that plant-derived biostimulants (PDB) and PGPR combinations have better biostimulation ef-fect compared to inoculation with microorganisms’ or the same type biostimulants com-binations. The protein hydrolysate in (BJ+AP+PH) combination may have functioned not only as phytohormones and amino acids source for soybeans, but also as an additional carbon and nitrogen source for bacteria vital processes [63], [64], especially under post-stress period in which nutrient exchange between the plant host and associated or symbiotic bacteria may have been negatively affected [65], [66]. The protein hydrolysate alone showed significantly better results, compared to the control in parameters such av-erage plant height, convex hull circumference, digital biomass, NDVI, PSRI and hue aver-age in early plant growth stage before stress induction, however, these results were insig-nificant compared to other biostimulants treatments. After stress induction period, no sig-nificant differences were obtained between protein hydrolysate and control variants. Seed coating with protein hydrolysate alone showed positive results in early soybean develop-ment stage, however, more research should be performed to investigate additional appli-cations, the most optimal application time and dose [67], which can be important factors for further plant development stages and stress response. Our result implies that protein hydrolysate could be perfect option to biostimulate plants in early growth stages, but for abiotic stress reduction it should be used additionally with other biostimulants, preferably with microbial products.

A. pascens are widely used in industry for organic compounds such d-malate and d-citramalate and enzymes such inulin fructotransferase and choline oxidase pro-duction [68], [69], [70], however, this bacterium potential in agriculture as a biostimulant is less studied. We found no scientific information about this species application on crop plants, only one investigation was published where A. pascens BUAYN-122 strain effi-ciency was analysed on lettuce and celery in hydroponic cultivation. In that research A. pascens BUAYN-122 strain significantly promoted the main root length, lateral root num-ber and fresh weight of lettuce and celery seedlings. Compared with the control group, the fresh weight of lettuce and celery inoculated with A. pascens BUAYN-122 increased by 144.4% and 300.9%, respectively [71]. Our isolated A. pascens 13LEP5 strain used in com-binations demonstrated strong biostimulation effect before (VC growth stage) and after stress induction (V3 growth stage). Especially, strong biostimulation were achieved in post stress period, where single inoculation of A. pascens 13LEP5 strain was applied. Soybean average height and 3D leaf area were 45 % and 92 % significantly higher compared to the control plants, respectively. These results could be associated with A. pascens ability to produce auxin indole-3-acetic acid (IAA) which is important phytohormone for plant cell division, elongation and differentiation [50], [72]. IAA also plays a crucial function in ex-panding plant root cell walls, resulting better water absorption in drought conditions and substantial enhancement in root exudation. Higher exudates extraction influences the ac-tivity and composition of rhizosphere bacteria [72]. Different studies show that IAA im-proves drought tolerance by increasing sugar and proline contents, activating auxin, ab-scisic acid and jasmonic acid related genes and inhibiting senescence genes [73], [74]. Some Arthrobacter species have a wide range of different abiotic stress tolerance mecha-nisms, such as the expression of ACC deaminase that reduces stress-induced ethylene lev-els, and the build-up of osmolytes such as glycine-betaine [75]. So, A. pascens 13LEP5 strain showing high potential as a plant biostimulant to increase plant growth and development and reduce abiotic stress, especially by incorporating this strain into combination with B. japonicum and protein hydrolysate.

Our research showed that biostimulants have stronger influence on plant spectral parameters and pigments accumulation before stress induction, compared to the efficiency in post stress period. NDVI, which measure vegetation health and greenness [76], analysis conducted before stress induction showed that all soybean seedlings treated with biostim-ulants exhibited significantly higher photosynthetic activity compared to the control group. NPCI shows pigments and chlorophyll content [77] and PSRI measures plants stress and senescence [78], these values range from −0.1 to 0.2 and −0.1 to 0, respectively and show healthy vegetation, these values increase in response to abiotic stress (i.e., heat nutrient limitation) [79], [80]. NPCI demonstrated that there was no pigments and chloro-phyll deficiency in any of the plants analysed, PSRI revealed that all plants experienced minimal stress and senescence, but stress and senescence levels were significantly higher in the control group compared with the groups where biostimulants were used. Hue Av-erage, which demonstrates leaf colour shade and is related with leaf chlorophyll content [81], showed that soybean seedlings treated with biostimulants, except (BJ+AP+PH) treat-ment, were significantly greener than control plants. Saturation Average, which shows colour brightness [82], demonstrated that all soybean seedlings were in normal physio-logical state and had quite intensive leaf saturation, only (BJ+AP+PH) demonstrated sig-nificantly better result compared to the control. After stress induction and recovery period no significant NDVI difference were founded between examined plants. All soybean plants showed a slight decrease in chlorophyll content, however, the differences were not statistically significant, and chlorophyll levels remained within the normal range. PSRI values slightly increased after stress, indicating a small stress increase, however, differ-ences between values obtained were statistically insignificant and levels of stress and se-nescence were still low. Plants treated with (BJ+AP+PH) demonstrated significantly more intensive saturation compared to the control, however, all plants physiological state, based on saturation intensity, was normal. Although available information about the effect of biostimulants on the values of crop vegetation indices is limited, some studies shows that biostimulation application can increase different vegetation indexes. Campobenedett with team found that NDVI of tomato plants was increased after tannin-based biostimu-lants treatment [83]. Xu, Chenping, and Beiquan Mou found that fish-derived protein hy-drolysate significantly enhanced lettuce chlorophyll content, photosynthetic rate, stomatal conductance [84]. Agliassa and her colleges demonstrated that plant protein hydrolysate increased chlorophyll level and promoted a faster and more efficient Capsicum annuum re-covery after drought [85]. Mateus Neri Oliveira Reis and his team showed that microbial inoculation with PGPR significantly improved soybean photosynthetic rate and chloro-phyll index [86]. Only Guillard, indicated that biostimulants containing seaweed extract have not increased NDVI under less than extreme heat-stress conditions, water-stress conditions, or both [87]. Based on out and other authors findings it could be that our in-duced stress was not severe enough or was too short to cause chlorophyll degradation, and to see differences between biostimulants efficiency on abiotic stress reduction. It also may indicate that the tested biostimulants were more effective in optimizing prestress physiological condition than in maintaining differential responses during the post stress period. However, among the tested biostimulators treatments, the (BJ+AP+PH) triple com-bination demonstrated partial retention of its beneficial effect, particularly in colour relat-ed parameters, suggesting a potential synergistic interaction and potential on abiotic stress reduction.

Statistical analysis demonstrated that biometric parameters were more responsive to biostimulants application than pigment-related spectral indices. Significant differences among control and treated plants persisted in biometric parameters even after stress in-duction period, particularly for leaf area, plant height, biomass accumulation and canopy architecture. In contrast, pigments associated indices (NDVI, PSRI, Hue) lost statistical significance after stress induction, while NPCI showed no significant differences throughout investigation period. These findings revealed that the tested biostimulants demonstrated a more stable long-term effect on structural plant development than on pig-ment accumulation. Md Abdul Mannan and his colleagues demonstrated that water defi-cits reduced chlorophyll pigments in soybean leaves under drought conditions [88]. Under drought stress, plants induce stomatal closure to prevent water evaporation, stomatal clo-sure limits the diffusion of CO2 into the leaf mesophyll and subsequently restricts the rate of photosynthesis [89]. Low temperatures also have negative impact on photosynthesis, can slow down metabolic processes and reduce photosynthetic enzymes responsible for chloroplast function [89]. Our results indicating that the physiological advantages of B. japonicum, A. pascens and protein hydrolysate are probably based on an enhanced nutrient uptake performance and hormonal control of growth, instead of a major regulation of the processes important for leaf pigments formation. We also found plant canopy architecture reorganisation after stress induction in variants where boiostimulant were applied. Soy-bean plants treated with biostimulants has not only enhanced biomass accumulation and canopy development but also increased canopy light penetration and plant height. It sug-gests that biostimulants modified canopy architecture rather than simply increasing can-opy density. This plant structural reorganisation reduced self-shading and improved in-ternal light distribution.

## 5. Conclusions

In this research we found that biostimulants application on seed has positive effect on soyabean biometric parameters in early plant development stage and post stress peri-ods. However, more stable long-term effect was found on structural plant development parameters, than on pigment accumulation. After abiotic stress induction the best results on plant biometric parameters were found where (AP) and (BJ+AP+PH) combination was inoculated. (BJ+AP+PH) combination was the only effective treatment, which showed significantly different results in pigments indices, compared to the control, after stress period. Based on our results (AP) and (BJ+AP+PH) have strong biostimulatation efficiency and hight potential on abiotic stress reduction. However, field trials must be performed to ob-tain the appropriate time, dosage and necessaryapplication number of these biostimula-tors. It is also important to evaluate (AP) and (BJ+AP+PH) stability and consortium com-patibility for long term storage as a product.

## Supporting information 66

No supplementary material

## Acknowledgments

This research was funded from the Research Council of Lithuania (LMT) under agreement no. S-AGROECOLOGY-25-1, as a part of EU Agroecology partnership project CoolFarmLab (Cool-FarmLab CO-creation Of Pathways to foster agroecoLogical transition of FARMs driven by Living LAB approach).

